# Reference trait analysis reveals correlations between gene expression and quantitative traits in disjoint samples

**DOI:** 10.1101/489542

**Authors:** Daniel A. Skelly, Narayanan Raghupathy, Raymond F. Robledo, Joel H. Graber, Elissa J. Chesler

## Abstract

Systems genetic analysis of complex traits involves the integrated analysis of genetic, genomic, and disease related measures. However, these data are often collected separately across multiple study populations, rendering direct correlation of molecular features to complex traits impossible. Recent transcriptome-wide association studies (TWAS) have harnessed gene expression quantitative trait loci (eQTL) to associate unmeasured gene expression with a complex trait in genotyped individuals, but this approach relies primarily on strong eQTLs. We propose a simple and powerful alternative strategy for correlating independently obtained sets of complex traits and molecular features. In contrast to TWAS, our approach gains precision by correlating complex traits through a common set of continuous phenotypes instead of genetic predictors, and can identify transcript-trait correlations for which the regulation is not genetic. In our approach, a set of multiple quantitative “reference” traits is measured across all individuals, while measures of the complex trait of interest and transcriptional profiles are obtained in disjoint sub-samples. A conventional multivariate statistical method, canonical correlation analysis, is used to relate the reference traits and traits of interest in order to identify gene expression correlates. We evaluate power and sample size requirements of this methodology, as well as performance relative to other methods, via extensive simulation and analysis of a behavioral genetics experiment in 258 Diversity Outbred mice involving two independent sets of anxiety-related behaviors and hippocampal gene expression. After splitting the dataset and hiding one set of anxiety-related traits in half the samples, we identified transcripts correlated with the hidden traits using the other set of anxiety-related traits and exploiting the highest canonical correlation (*R* = 0.69) between the trait datasets. We demonstrate that this approach outperforms TWAS in identifying associated transcripts. Together, these results demonstrate the validity, reliability, and power of the reference trait method for identifying relations between complex traits and their molecular substrates.

**AUTHOR SUMMARY:** Systems genetics exploits natural genetic variation and high-throughput measurements of molecular intermediates to dissect genetic contributions to complex traits. An important goal of this strategy is to correlate molecular features, such as transcript or protein abundance, with complex traits. For practical, technical, or financial reasons, it may be impossible to measure complex traits and molecular intermediates on the same individuals. Instead, in some cases these two sets of traits may be measured on independent cohorts. We outline a method, reference trait analysis, for identifying molecular correlates of complex traits in this scenario. We show that our method powerfully identifies complex trait correlates across a wide range of parameters that are biologically plausible and experimentally practical. Furthermore, we show that reference trait analysis can identify transcripts correlated to a complex trait more accurately than approaches such as TWAS that use genetic variation to predict gene expression. Reference trait analysis will contribute to furthering our understanding of variation in complex traits by identifying molecular correlates of complex traits that are measured in different individuals.

## INTRODUCTION

A major goal of complex trait analysis is to discover pathways and mechanisms associated with disease. By definition, these traits exhibit hallmarks of genetic complexity including pleiotropy, epistasis, and gene-environment interaction. Genetic mapping is a powerful approach for detecting quantitative trait loci that influence complex trait variation, but it has limited power for detecting small effect loci and can suffer from poor mapping resolution, hindering the identification of causal genes. Moreover, these causal genetic variants do not always reside in relevant therapeutic targets. Therefore, many systems genetic strategies have emerged to correlate complex traits directly with molecular phenotypic variation, with the goal of constructing molecular networks that are correlated with trait variation from a trait-relevant tissue or cell population.

Ideally, trait correlation networks are constructed using direct phenotypic measurements for each member of a population. However, there are wide-ranging questions for which this approach is infeasible or impossible because it is physically, technically, or financially impossible to obtain all of the measures in the same individuals. To refer to phenotypes whose measurement on the same individual is infeasible or impossible, we will use the term incompatible phenotypes. Incompatible phenotypes arise in common experimental designs such as studies of susceptibility to exposure effects where the exposure affects physiology (e.g. predisposition to psychostimulant addiction) or studies of disease that relate early stage changes to late stage outcomes (e.g. early molecular correlates predictive of Alzheimer’s disease risk). Moreover, incompatible phenotypes arise when the original study population no longer is available but there is a desire to extend the study to a new set of traits, a situation that is common in human genetic analyses. Finally, phenotypes could be incompatible for strictly financial or logistical reasons, for example due to prohibitively high costs of genomic assays in large cohorts, leading to fractional collection of data on some samples and more thorough characterization of others.

One emerging approach for relating gene expression and complex traits measured in different cohorts of genetically diverse individuals is to exploit genetic variants that affect gene expression (eQTL) to impute transcript abundance from genotypes alone (Gamazon *et al*. 2015; Gusev *et al*. 2016a; b; Mancuso *et al*. 2017; Barbeira *et al*. 2017). This enables estimation of the association between imputed gene expression and complex traits, an approach that has been called a transcriptome-wide association study (TWAS; Gusev *et al*. 2016a). However, the TWAS approach suffers from several limitations, most notably a reliance on strong local (presumably *cis*-acting) eQTL and consequent inability to impute transcript abundance for genes without detected eQTL. In contrast to using sparse, discrete *genotypes* to impute per-individual gene expression and infer correlation to complex traits, our approach uses shared variation across a rich set of quantitative, multidimensional *phenotypes* to infer gene expression correlates of phenotypic variability.

Rather than impute gene expression from genetic data, another strategy is to impute phenotypic data from other phenotypes. Hormozdiari et al. (2016a) used this approach to impute unmeasured phenotypes in the context of genome-wide association studies (GWAS; Hormozdiari *et al*. 2016a). Specifically, the method of Hormozdiari et al. (2016a) uses the correlation structure in one set of traits to predict a single unmeasured target trait in a second cohort using only phenotypic data. In the present study, we extend this strategy to multivariate phenotyping and apply it to transcriptomics, providing a precise transcript-to-trait correlation approach that can be compared to the TWAS method.

We outline a simple method, reference trait analysis, to study relations between a set of complex traits of interest (*target traits*) and a set of high-dimensional molecular traits obtained in disjoint subsets of individuals. Reference trait analysis relates these two incompatible, multidimensional sets of phenotypes indirectly through the use of a shared set of *reference traits* measured in all individuals. Since target and molecular traits are not measured in the same individuals, direct comparisons are impossible. Instead, we relate these traits through reference traits. Reference traits are best chosen with *a priori* knowledge that they share biological underpinnings with target traits. This relationship between reference and target traits is exploited to compute scores from reference traits that capture variation in unmeasured target traits and can be directly related to transcriptional profiles. By design, our method is robust to the detection of transcript-trait associations for which the regulation is not genetic or is characterized by multiple weak, indirect genetic effects. Therefore, it captures biological variability associated with both genetic and environmental sources of vulnerability, and has the potential to identify molecular networks of complex trait variation even when there is insufficient power to detect a quantitative trait locus or genome-wide significant SNP association.

In this study we develop and evaluate the reference trait analysis method using data from a previously published behavioral study of Diversity Outbred mice (Logan *et al*. 2013). Diversity Outbred mice are genetically unique; consequently, per subject terminal traits such as brain gene expression can only be obtained in a single exposure condition. However, the approach we propose can be useful in any heterogeneous population for which a common reference set of traits is assessed. Our assessment data set consists of multiple measures of anxiety-related traits in a sample of Diversity Outbred mice, all of whom have been subjected to brain transcriptional profiling as well as measurements of two sets of related behaviors. We present an overview of our method, use these data to assess sample size requirements, and quantify the method’s reliability across a range of target-reference trait correlations. Finally, we test whether the reference trait method more faithfully recovers trait-gene expression correlations than the TWAS approach.

## RESULTS AND DISCUSSION

### Outline of Approach

The reference trait analysis procedure is straightforward, and relies on well-characterized canonical correlation analysis. Beginning with a population of individuals, reference traits (labeled using the variable *U*) are measured on all individuals, target traits (labeled with *V*) on the *training* cohort, and high dimensional molecular traits (labeled with *M*) on the *testing* cohort (Figure 1). Although target traits and their molecular correlates are of primary interest, the choice of reference traits is an important aspect of the method. First, as we will show, the strength of the multivariate relationship between target and reference traits is a key parameter determining the power to detect trait-transcript correlations. Second, because our method leverages shared variation between target and reference traits, it identifies trait-transcript correlations driven by the portion of target trait variation that is shared with reference traits. For example, studying addiction-related traits using novelty behaviors as reference traits would be expected to uncover transcripts associated with addiction behaviors through biological pathways that also contribute to the etiology of novelty-seeking behaviors.

**Figure 1:**
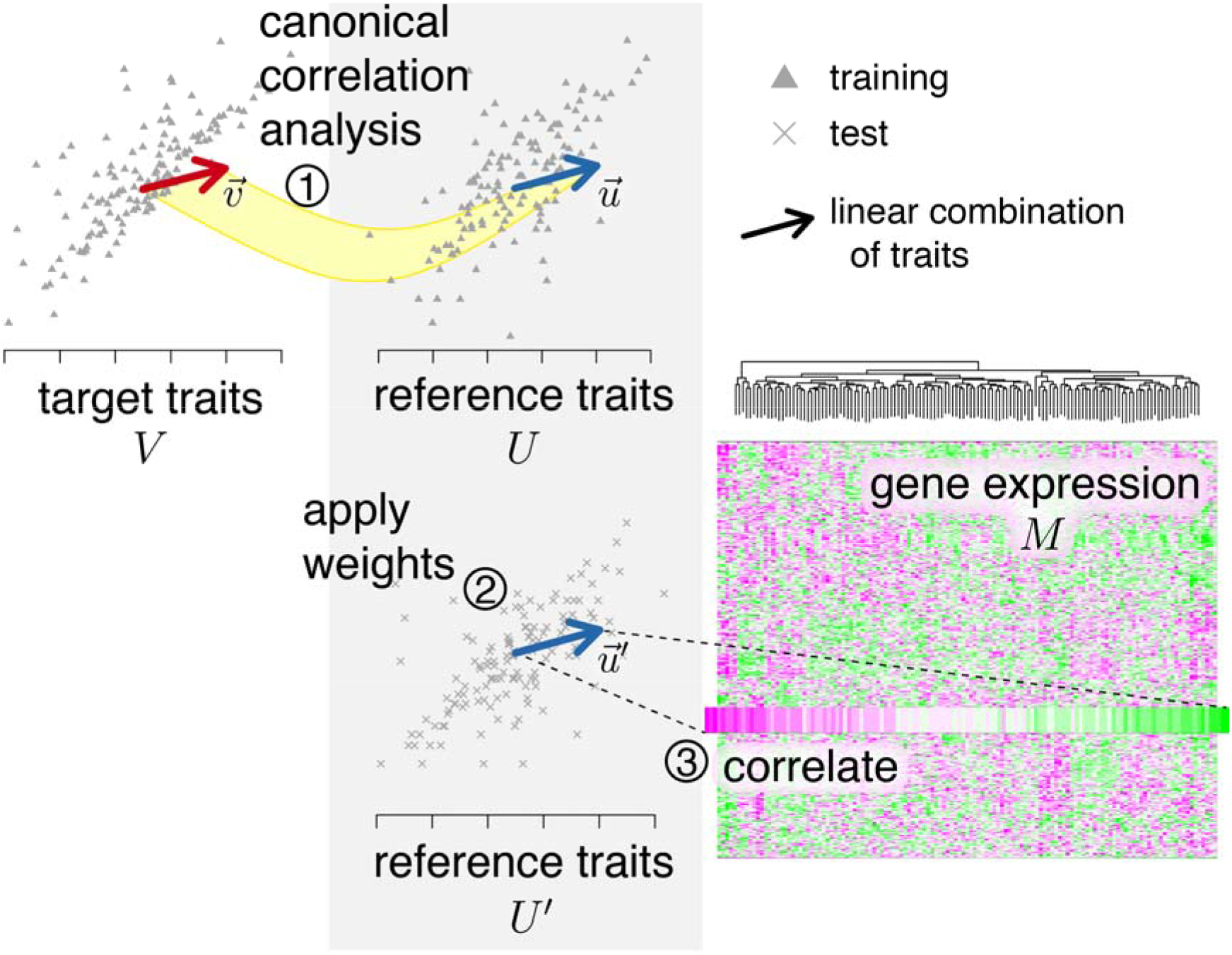
Overview of reference trait analysis. Target and reference traits are measured in a set of training individuals (top plots; grey triangles), while reference traits and gene expression are measured in test individuals (bottom plots and X symbols). (1) Canonical correlation is used to identify a linear combination of reference traits (top blue arrow) that best captures variation in the traits of interest (red arrow; yellow curve connecting arrows represents canonical correlation analysis). (2) The weights derived from canonical correlation analysis are applied to reference traits in the testing population to derive reference trait scores for each individual (projected reference traits; bottom blue arrow). (3) Projected reference traits are correlated with molecular phenotypes.

To conduct reference trait analysis, we employ canonical correlation (Hotelling 1936), which can be thought of as a parent analysis of the more familiar multiple regression. A multiple regression of *Y* on *X* models the relationships between multiple *X* measures *X*_1_, *X*_2_,…, *X_p_* and univariate *Y*. In contrast, canonical correlation reveals the magnitude and nature of relationships between multivariate *U* and *V*, e.g. *U*_1_, *U*_2_,…, *U_p_* and *V*_1_, *V*_2_,…, *V_q_*. Specifically, canonical correlation identifies linear combinations of two multivariate measures *U* and *V* such that the (univariate) linear combinations of each measure 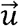 and 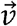, known as canonical variables, are maximally correlated. In this study we use canonical correlation to build linear combinations of reference traits (transforming *U* to 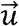) that maximize shared variance with target traits (*V*) in the set of training individuals. The possible number of canonical variables is limited to the size of the smaller of *U* and *V*, and each successive covariate captures a diminishing proportion of the shared variance between the traits. In this study we focus on the first canonical variable, 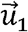 or 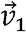, which explains the largest fraction of shared variance between *U* and *V*. This quantity can be thought of as a summary of each set of traits analogous to their first principal component, but rather than being aligned with the axis of maximal variation *among* a single set of variables, it is aligned in the direction of maximal shared variation *between* the two sets of traits *U* and *V*. For datasets with a very large number of reference and/or target traits (i.e. *p* ≫ *n*), sparse canonical correlation analysis (Witten and Tibshirani 2009; Wilms and Croux 2016) may reduce over-fitting, but this situation is not common when relating two sets of traits *U* and *V* that contain organism-level phenotypes as opposed to molecular features.

The analysis of training data defines canonical coefficients that can be used to compute first canonical variables from individual-level trait data (i.e. transform *U* to 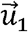 or *V* to 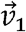). We use these coefficients learned from the training data (Figure 1, top) to transform reference trait data from the testing cohort *U*’, which projects these data in the direction of maximal shared variation with target traits. Thus, these “projected” traits 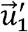 optimally capture the portion of variation shared between reference and target traits due to their underlying genetic and environmental covariation. Projected traits are then compared to high-dimensional genomic measurements to extract molecular phenotypes in one sample set that co-vary with target traits from another group (Figure 1, bottom right).

### Transitive reliability captures global patterns of covariation between incompatible traits

Reference trait analysis reveals covariation between molecular phenotypes and target trait variation. There are many possible applications of this strategy. For example, in addiction research, many studies evaluate transcriptional response to drug exposure but are unable to evaluate the predisposing characteristics of a drug naïve brain that associate with addiction-related behaviors. Using a reference trait strategy, one can evaluate the transcriptomes of drug naïve brains and relate them to the response to drug self-administration through a set of reference traits that do not involve drug exposure. We have previously estimated the association of novelty seeking and drug self-administration in mice, revealing a canonical correlation of 0.61 among these sets of traits (Dickson *et al*. 2015).

To evaluate whether the reference traits strategy could be applied to find transcriptional correlates of drug self-administration, we used a dataset where reference, target, and molecular trait profiling were performed on the same individuals to allow for assessment of the accuracy and robustness of the method. In this data set, transcriptional profiles, target traits, and reference traits are available for all individuals. This allows evaluation of the properties of the reference trait strategy, including robustness and sample size requirements. Specifically, we studied relationships between two distinct sets of anxiety-related traits and hippocampal gene expression, where all traits were measured in each of *N* = 258 Diversity Outbred mice (Logan *et al*. 2013). The anxiety-related traits consisted of eleven measurements of open-field arena exploration behaviors and five measurements of light-dark box behaviors (Supplementary Table 1). A canonical correlation analysis of these two sets of traits yielded a statistically significant model (F_55,1123.75_ = 4.48, *p* < 2×10^-16^, Wilk’s λ = 0.400) that had a first canonical correlation coefficient of magnitude 0.69. This was higher than all univariate correlations between open-field and light-dark box traits (median 0.11, maximum 0.65), and similar in magnitude to the shared variation revealed by the first canonical variable in the motivating analysis of novelty-related behaviors and cocaine self-administration (Dickson *et al*. 2015). We arbitrarily designated the open-field traits as target traits and light-dark box traits as reference traits. For reference trait analysis, we hid gene expression data for some mice (training set) and open-field data for the remaining mice (testing set).

In this evaluation of reference trait analysis, we know the true values of all hidden data and can directly evaluate the power of the method to reveal gene expression patterns associated with target trait variation. Specifically, we estimate canonical coefficients (weights to calculate canonical variables) from the training set and use them to calculate projected traits 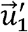 in the testing set. To quantify the performance of reference trait analysis when the true answer is known, we computed correlations in testing set animals between gene expression *E* and either (1) the first projected trait 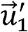, cor (*E*, 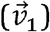) or (2) the first canonical variable computed using hidden target traits 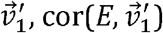. The latter quantity, the “truth”, is unavailable in a real application of reference trait analysis. A set of reference traits that perfectly captures all variation in target traits would result in a vector of gene expression-trait correlations that is identical whether the target traits were known or projected from reference traits (i.e. the reference traits serve as a perfect surrogate for target traits). We define *transitive reliability* as the correlation between these vectors i.e. 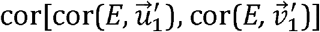. High transitive reliability would indicate that strong correlations between gene expression and target traits are likely to be identified using projected traits.

Transitive reliability, estimated using real gene expression data and simulated canonical variables with known correlation, scales linearly with the magnitude of the canonical correlation coefficient (Figure 2A), confirming our intuition that greater sharing of variation between target and reference traits increases the utility of leveraging reference traits to understand target trait variation. We divided the anxiety dataset into equally sized subsets (partially overlapping for larger sample sizes) to examine the dependence of transitive reliability on sample size. The canonical correlation was upwardly biased for small sample sizes (*N* < 90; data not shown), as has previously been recognized (e.g. Thompson 1990). When we used Wherry’s correction as suggested by Thompson (1990), canonical correlations no longer depended on sample size (linear model; *p* > 0.8). Overall, transitive reliability asymptotically approached the magnitude of the canonical correlation coefficient calculated from the full dataset (Figure 2B, black line), demonstrating that global patterns of trait-gene expression correlation can be recovered with relatively modest sample sizes using the reference trait approach. In contrast, weights from the smallest (fifth) canonical covariate, which captures little shared variation between datasets, produced low transitive reliabilities (median 0.11).

**Figure 2:**
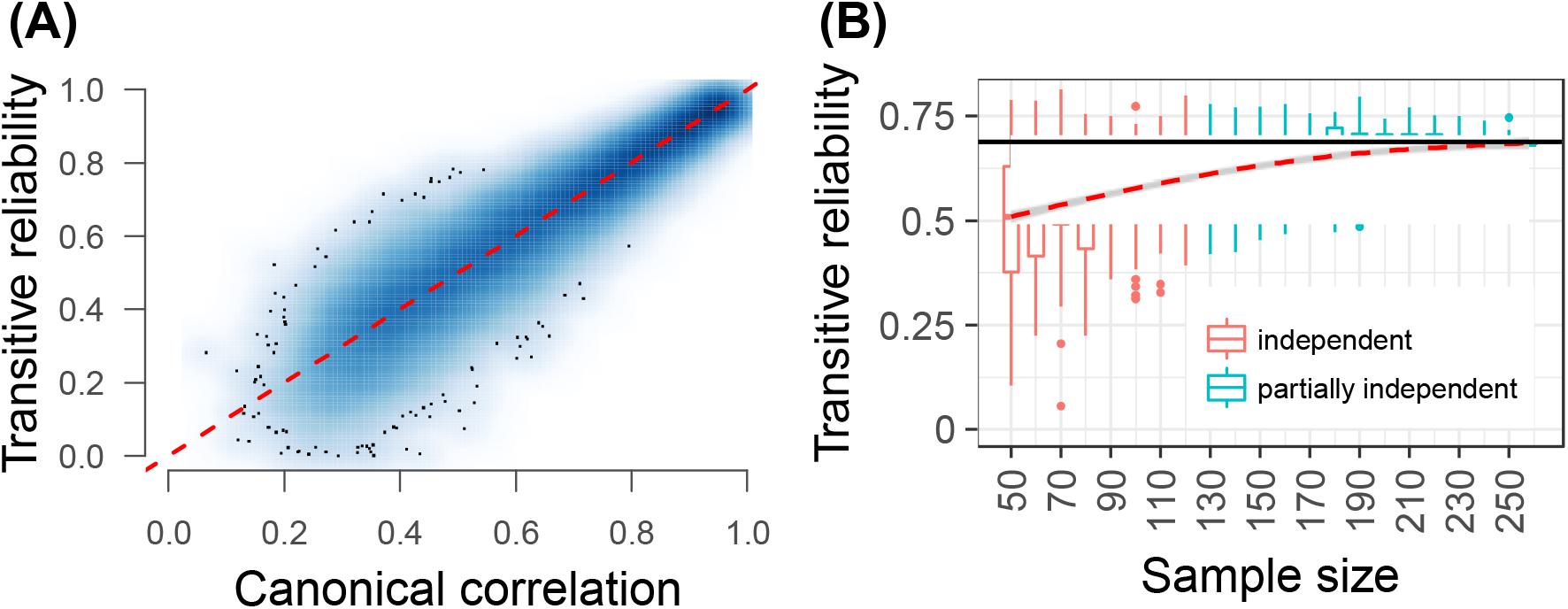
Reference trait analysis reveals overall patterns of covariation between incompatible traits. (A) Relationship between canonical correlation and transitive reliability. To evaluate the mathematical relationship between these quantities, we simulated two vectors with known correlation to represent the canonical covariates, and calculated transitive reliability with real gene expression data. Canonical correlation shown is absolute value, and transitive reliability is sign-matched. (B) Sample size increases lead to higher and more precise transitive reliability. Plot shows transitive reliability estimated using anxiety data with animals subsampled as described in the main text. Sample size on *x*-axis indicates the number of individuals used in each of the training and testing groups (the number of individuals phenotyped for target traits and the number with high-dimensional molecular phenotypes, respectively). Black line indicates magnitude of first canonical correlation calculated from full dataset. Color indicates whether training and testing groups were fully or partially independent.

### Reference trait analysis successfully identifies known trait correlations

Ultimately, the primary goal of reference trait analysis is to identify molecular correlates of unmeasured phenotypes. To discover these correlates, individual gene expression levels are correlated to projected traits. To test this strategy, we first employed reference trait analysis on the anxiety-related phenotype data described above. After randomly splitting the dataset and withholding open-field data (arbitrarily designated as target traits) in half the individuals, we identified gene expression levels correlated to projected reference traits. We found high overlap between the genes most strongly correlated with hidden target trait canonical variable 1 and those most strongly correlated with projected traits (23% overlap among genes with top 5% of correlations to each trait, compare to 2.5% expected overlap; *p* < 1×10^-15^, Fisher’s Exact Test). Across all genes, including those with weaker correlations, we found that the vector of trait-gene expression correlations computed using reference trait analysis showed significant similarity to the true correlations (*p* < 0.001, permutation test using generalized Jaccard similarity statistic). Moreover, in contrast to the alternative methods for identifying trait-gene expression correlations discussed above, some correlations detected using reference trait analysis involved genes with no significant eQTL (e.g. 42% of top 50 correlations). These genes, which are demonstrably associated with trait variation, would not be detectable using TWAS type approaches.

To examine the power and robustness of reference trait analysis across a wide range of biologically plausible parameter values, we conducted extensive simulations. We simulated data across a range of sample sizes (100, 200, 300,…, 1000,1200,1400,…, 2000) and enforced a similar covariance structure to the observed data. Specifically, data were simulated using observed covariances within each set of anxiety traits, but we perturbed covariances between the two sets of traits in order to generate datasets with varying canonical correlations. We then simulated gene expression levels with known correlation to the first target trait canonical variable, 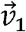 (ρ = 0.2, 0.225, 0.25,…, 0.9 with 20 genes each). We simulated trait data and gene expression data at random for each of 1,000 simulations for each sample size.

For each simulation, after hiding target traits in half the individuals and gene expression data in the other half, we conducted reference trait analysis. We computed projected reference traits, correlated to gene expression, and quantified performance as the fraction of true trait-gene expression correlations that were detected using a 10% false discovery rate (FDR) threshold. For high trait-gene correlations (ρ > 0.6) and strong target-reference trait canonical correlations (R = 0.7 or 0.9), the correlation of interest was essentially always detected (Figure 3). For lower target-reference trait canonical correlations (R = 0.5), even relatively modest true trait-gene expression correlations (e.g. ρ = 0.3) were often detected with sample sizes above ~300 individuals (Figure 3). Thus, reference trait analysis was a highly effective means for identifying trait-gene expression correlations across a diverse range of practical sample sizes, typical values for trait-to-gene expression correlation, and canonical correlation parameters.

**Figure 3:**
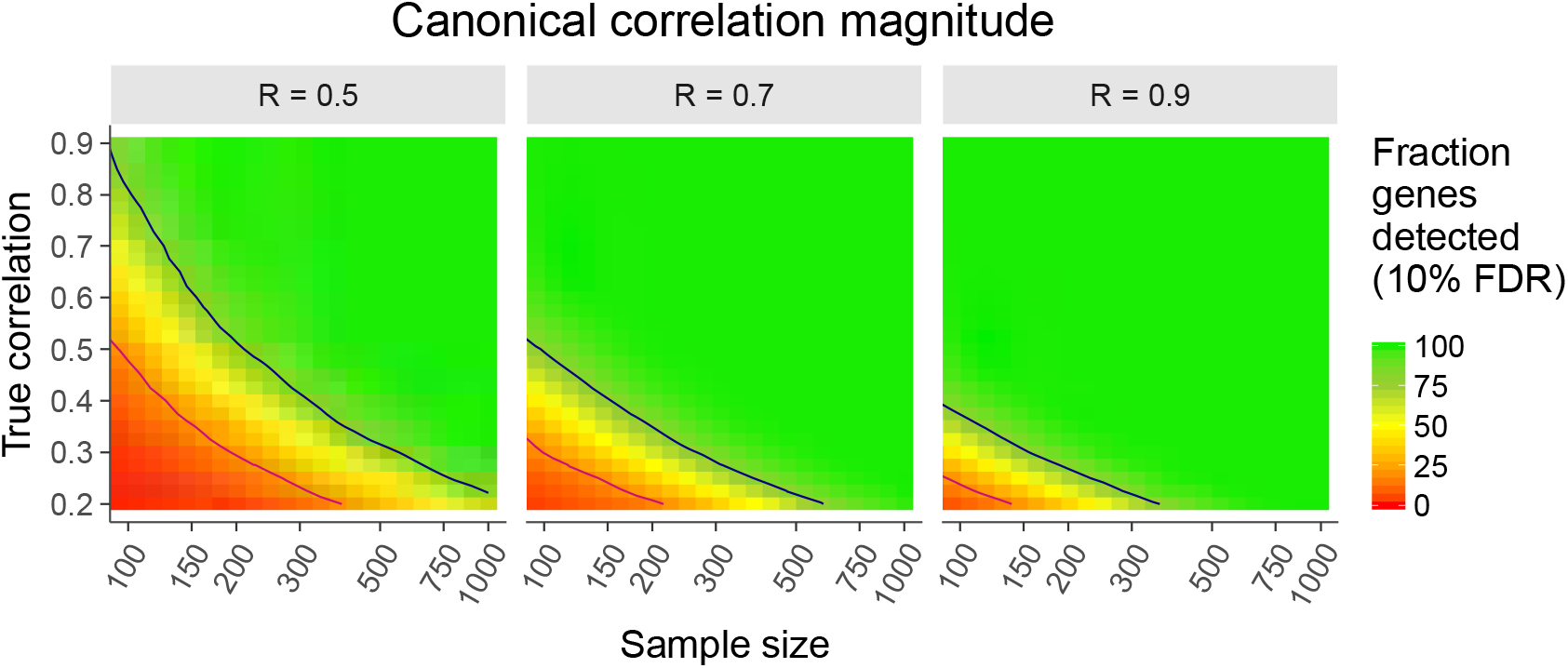
Reference trait analysis identifies simulated trait-gene expression correlations across a wide variety of parameter values. Sample size plotted along *x-* axis is the number of individuals used in each of the training and testing groups (equal sample size for the two groups, where the training group consists of individuals phenotyped for target traits and the testing group those with high-dimensional molecular phenotypes). True correlation (*y*-axis) indicates correlation between first target trait canonical variable 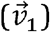 and simulated gene expression. Facets indicate magnitude of canonical correlation coefficient between reference and target traits (*R* listed along grey strips, ±0.02). Navy and magenta contour lines depict regions above/below which trait-gene expression correlations are detected >80% and <20% of the time, respectively.

### Comparison of reference trait analysis to related approaches

An alternative approach to identifying genes associated with complex traits is to make use of known genetic variation that regulates gene expression (gene expression QTL or eQTL). There has been considerable recent interest in methods that integrate complex trait associations and gene expression genetics in order to identify genes whose expression is associated with trait variation (Nica *et al*. 2010; Wallace *et al*. 2012; He *et al*. 2013; Gamazon *et al*. 2015; Gusev *et al*. 2016a; Zhu *et al*. 2016; Hormozdiari *et al*. 2016b; Wen *et al*. 2017; Hauberg *et al*. 2017). Several methods perform tests of the hypothesis that genome-wide association (GWA) signals and eQTLs are truly colocalized versus independent but appearing colocalized due to linkage disequilibrium (Nica *et al*. 2010; Wallace *et al*. 2012; Giambartolomei *et al*. 2014; Fortune *et al*. 2015; Zhu *et al*. 2016; Hormozdiari *et al*. 2016b; Wen *et al*. 2017; Hauberg *et al*. 2017). Another approach that is more directly applicable to the experimental designs studied herein is to harness strong genetic predictors of gene expression variation (eQTL) to impute transcriptomes in genotyped and phenotyped cohorts, which allows detection of trait-expression correlations (the TWAS approach; Gamazon *et al*. 2015; Gusev *et al*. 2016a; b; Mancuso *et al*. 2017; Barbeira *et al*. 2017). TWAS is an approach that is complementary to reference trait analysis, and has been a particularly powerful method for discovery of candidate genes driving GWA signals detected in very large human cohorts (tens or hundreds of thousands of individuals). Supplementary Figure 1 provides a comparison of genotype, phenotype, and gene expression data in the reference traits and TWAS strategies. One weakness of the TWAS approach is that it hinges on the presence of detectable eQTL (typically local, presumably cis-acting eQTL; but see He *et al*. 2013; Vervier and Michaelson 2016). In humans, even panels of 1,000 individuals with gene expression measurements only result in a modest number of genes (500-4,000) with significant *cis*-heritability that can be imputed in the cohort lacking gene expression data (Gusev *et al*. 2016a). In contrast, reference trait analysis has no requirement for detection of eQTLs, and therefore it is amenable to detect of correlation of transcripts with complex expression regulatory mechanisms to traits of similarly complex regulation, and retains performance across lower sample sizes, as we demonstrate below.

Although TWAS and reference trait analysis utilize different data types, both are tools inferring relations between complex traits and transcript abundance, so we sought to compare their performance on the same dataset. For TWAS, we used methods implemented in the software suite PrediXcan (Gamazon *et al*. 2015). We randomly divided our anxiety dataset in half and considered open-field measurements as target traits. We withheld gene expression measurements in half the animals; therefore, only genotype and reference trait data were visible for all animals. We built predictive models of gene expression from the training cohort of mice, applied these models to impute gene expression in the testing cohort, and calculated correlations between imputed gene expression and a summary measure of the target traits (first canonical variable). We conducted 1,000 permutations with random 50:50 divisions of the anxiety dataset to account for stochastic sampling effects. For each replicate, we compared global trait-gene expression correlations for PredictDB-imputed gene expression versus those computed using projected traits obtained with our new method. In the former case, trait data is available and gene expression data is imputed, while in the latter case gene expression data is available and trait data is imputed.

For direct comparisons between reference trait analysis and TWAS, we ran reference trait analysis using only genes that were significantly predicted by the PredictDB module of PrediXcan (FDR < 5%; see Methods). Across the 1,000 permutations, we imputed gene expression for a mean 12,250 genes (range 11,640-12,750; mean represents ~70% of total 17,539 genes measured), indicating that a substantial fraction of genes has insufficient local genetic signal for accurate imputation. An advantage of reference trait analysis is that it is not limited by the presence of strong eQTL and all genes can be tested for association with projected reference traits. For each of the 1,000 permutations, we computed the transitive reliability of TWAS and of reference trait analysis. Reference trait analysis more accurately captured global patterns of trait-transcript correlation than TWAS (Figure 4). Specifically, transitive reliability for target trait first canonical covariate-gene expression correlations was higher using the reference trait approach (measured gene expression and projected reference traits) compared to the TWAS approach (imputed gene expression and measured traits) for 92.7% of simulations (Figure 4; Supplementary Figure 2 shows an example of results from one permutation). Thus, we show empirically that reference trait analysis outperforms TWAS in the mouse anxiety dataset.

**Figure 4:**
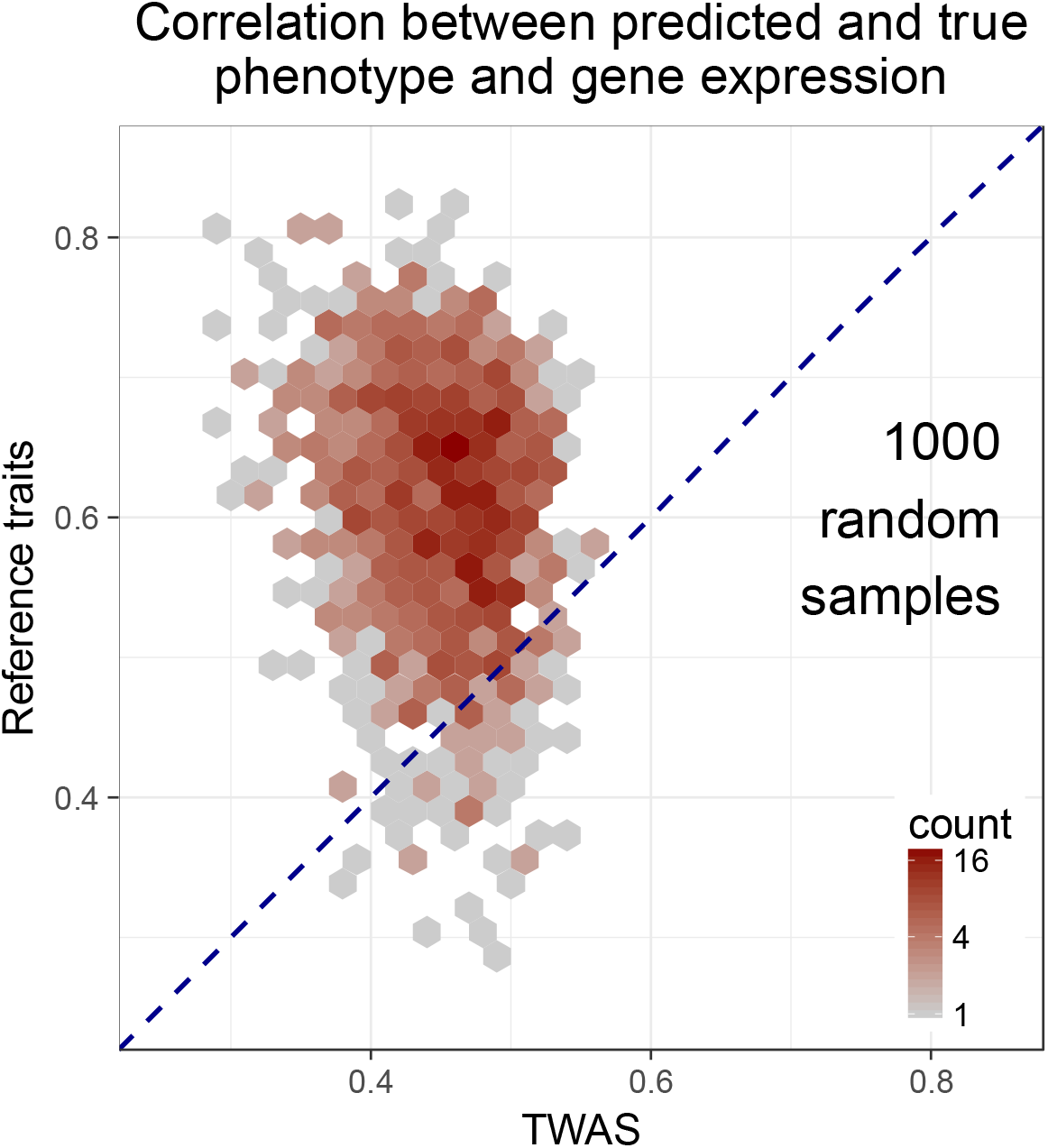
Reference trait analysis recovers true trait-gene expression correlations more accurately than TWAS. Binned hexagon plot shows the results of 1,000 random samples where the anxiety dataset was split into two halves randomly designated the training and testing groups. Reference trait analysis and TWAS were used to recover trait-gene expression correlations. The true values of both the trait and gene expression are known in this dataset, but were hidden when running reference trait analysis or TWAS. For each method, the correlation across all genes between predicted and true values was computed.

In addition to the quantitative comparison of the methods, we sought to determine which approach provided the best retrieval of known anxiety related genes. To perform this analysis we made use of GeneWeaver’s database of gene sets curated from multiple sources (Baker *et al* 2016). The top four hundred genes identified using each analysis method were entered as three gene lists, and each gene list was compared to every gene set in the GeneWeaver database via Jaccard similarity. For each, the top 249 similar gene sets were exported, and a rater with expertise in behavioral neuroscience who was blind to the analysis methods scored a combined list of all similar gene sets obtained in these three analyses. Gene sets were categorized discretely based on relevance to anxiety, with categories including irrelevant, generally relevant to brain or behavior, and specifically relevant to anxiety. We found that true open-field first canonical variable—gene expression correlations had highest relevance to anxiety. The top truly correlated genes were similar to gene sets more relevant to anxiety than those genes identified using reference traits or those using TWAS (*p* = 0.0065 and *p* = 1.5×10^-14^, respectively; two-sided Fisher’s Exact Test). Nevertheless, reference trait analysis performed significantly better than TWAS at identifying genes with similarity to anxiety-relevant gene sets (*p* = 7.3×10^-6^).

Finally, another alternative to relating traits and transcripts between population cohorts is to make use of polygenic risk predictors trained using genome-wide genotypes and phenotypes, and applied to individuals with genotypes but missing phenotypes (in this case, samples with only transcriptional profiles available) (Makowsky *et al*. 2011; Dudbridge 2013; Wray *et al*. 2013). However, theoretical considerations and empirical results suggest that this approach generally requires sample sizes much larger than 1,000 individuals to obtain accurate predictions (Dudbridge 2013). In the context of reference trait analysis, relating complex reference and target traits that share high canonical correlation implicitly leverages the common polygenic or omnigenic (Boyle *et al*. 2017) basis of these traits by making use of all of the information contained in continuous quantitative variation.

## Conclusions

We have described a general method for exploring trait covariation among incompatible and independently collected phenotypes studied in disjoint samples of genetically diverse individuals to extract molecular networks associated with disease. Our method utilizes canonical correlation analysis, a standard multivariate statistical method, to relate incompatible phenotypes using a set of reference traits measured on all individuals. Our analyses demonstrate that this approach performs well over a range of parameters typically encountered in the study of trait correlations, and under sample size requirements that are practical to obtain. This approach can be useful both for capturing global patterns of covariation between target traits and high-dimensional molecular phenotypes, as well as for identifying specific molecular correlates to target traits. Our method identifies trait-gene expression associations and we do not assert that these associations are necessarily causal, as has been recognized by studies relating GWAS results and eQTL (Gamazon *et al*. 2015; Gusev *et al*. 2016a; Hauberg *et al*. 2017).

When will reference trait analysis be a useful tool? Intuitively, and as demonstrated in Figure 3, large sample sizes, precise trait measurements, and high shared variance between reference and target traits would allow for the most accurate estimation of canonical correlation coefficients and high power to detect correlations to molecular phenotypes. Although our method could be applied in a wide variety of scenarios, it is likely to be particularly useful for studies of highly complex, polygenic, multidimensional traits (e.g. behavior, physiology, and morphology) in cohorts of modest size. As with any method that applies information learned from one cohort to biological measures from another cohort, reference trait analysis requires the absence of systematic differences (i.e. heterogeneity in population characteristics) between the training and testing cohorts. For very large cohorts of individuals where obtaining suitable reference traits may be difficult, polygenic scores based on either genetic predictors alone or on a combination of genetic and environmental risk factors (Dudbridge *et al*. 2017) may be a valuable approach for predicting phenotypic variation in a test cohort that can then be correlated with molecular networks.

Although our application of reference trait analysis involves correlations to high dimensional molecular phenotypes, the method could, in principle, be applied to any sets of phenotypes that are multivariate in nature. Moreover, the high relative performance of our method underscores the importance of extensive phenotyping using quantitative traits rather than relying on binary indicators of disease and disease-related phenotypes that may mask complex underlying etiologies. We anticipate that the framework outlined in this study will be increasingly useful as studies of diverse, genetically unique populations become more widespread. A useful future extension to this approach would incorporate statistical techniques such as sparse canonical correlation analysis (Witten and Tibshirani 2009; Wilms and Croux 2016), which could permit inference in phenome-level studies where the target or reference traits are high dimensional. Overall, our approach is likely to be particularly important in functional genomics studies, those utilizing post-mortem subjects, and large population studies in which individuals are unavailable for further characterization.

## Materials and Methods

### Mouse rearing and phenotyping

Diversity Outbred mice (J:DO, The Jackson Laboratory) are a heterogeneous stock derived from the same eight founder strains as the Collaborative Cross (Svenson *et al*. 2012; Churchill *et al*. 2012; Gatti *et al*. 2014; Chesler *et al*. 2016). In this study we used a subset (*N* = 258) of the 283 Diversity Outbred mice studied by Logan et al. (2013) with hippocampal gene expression profiled by RNA-Seq (see below). Mice in this study were from generations 4 to 5 (G4-G5) of the DO population. Briefly, each mouse was acclimated to the housing area, and subject to a brief testing battery which included a 20 minute novel open-field test and a 10 minute light-dark test, among other common behavioral tasks. The open-field and light-dark tests are used to measure exploratory activity and approach-avoidance behavior. Many complex trait measures can be extracted from these tasks. For this analysis, we chose two sets of informative measures (Supplementary Table 1). Complete details of animal rearing, husbandry and phenotyping are presented in Logan et al. (2013). Mice were sacrificed using decapitation which was necessary to preserve fresh brain tissue in the absence of drug or asphyxiation. All procedures and protocols were approved by The Jackson Laboratory Animal Care and Use Committee, and were conducted in compliance with the National Institutes of Health Guidelines for the Care and Use of Laboratory Animals.

### Genotyping

DNA was prepared from tail biopsies and samples were genotyped using the Mouse Univeral Genotyping Array (MUGA) (Morgan *et al*. 2016). We obtained genotypes at 7,802 markers from arrays processed by GeneSeek (Lincoln, NE). We used intensities from each array to infer the haplotype blocks in each individual DO genome using a hidden Markov model (Gatti *et al*. 2014).

### Gene expression profiling

Total hippocampal RNA was isolated using the TRIzol^®^ Plus RNA purification kit (Life Technologies Corp., Carlsbad, CA) with on-column DNase digestion. Samples for RNA-Seq analysis were prepared using the TruSeq kit (Illumina Inc., San Diego, CA) according to the manufacturer’s protocols and subjected to paired-end 100 base pair sequencing on the HiSeq 2000 (Illumina) per manufacturer’s recommendations. RNA sequencing was performed in nine sequencing runs with two technical replicates for each sample, resulting in an averaging sequencing depth of approximately 24 million reads per sample after pooling technical replicates. To obtain estimates of gene expression, we aligned reads to individualized diploid genomes using the bowtie aligner (Langmead *et al*. 2009) and quantified transcript abundance by allocating multi-mapping reads using the EM algorithm with RSEM (Li and Dewey 2011) as described in Munger et al. (2014). Raw counts in each sample were normalized to the upper quartile value and transformed to normal scores.

### Reference trait analysis

We conducted reference trait analysis using R version 3.3.2 (R Core Team 2016). Canonical correlation analysis was carried out using the cancor function in base R. We regressed out the effect of sex on each phenotype because it is not of primary interest in this study. An example walk-through of a reference trait analysis and code to carry out the analyses described in this paper are available at https://daskelly.github.io/reference_traits/reference_trait_analysis_walkthrough.html.

To examine the power and robustness of reference trait analysis, we simulated data with varying sample sizes and canonical correlation coefficients. We based our simulations on the anxiety phenotype data, consisting of open-field exploration and light-dark box behavioral measures. Specifically, for each of 1,000 simulations we started with the covariance matrix computed from five open-field and five light-dark box traits and randomly increased or decreased each of the 5×5=25 inter-dataset covariances by 20%. We then simulated multivariate normal phenotype data with the specified covariance matrix. This procedure resulted in two multivariate datasets (simulated open-field and light-dark box traits), where the covariance structure *within* each dataset was similar to that in the real data but with different covariances *between* datasets. When a canonical correlation analysis was carried out on each pair of simulated datasets, the magnitude of the first canonical correlation coefficient varied between *R* = 0.35 and *R* = 0.98, due to the variation in interdataset covariances.

We simulated gene expression traits with exact correlation to the first target trait canonical variable 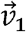 in the simulated dataset. In order to simulate a random vector of observations with defined correlation to an existing vector, we took advantage of the geometric property that the cosine between two mean-centered vectors equals their correlation. Therefore, a random vector with defined correlation to an existing vector can be computed by starting with random draws from a normal distribution, mean-centering, and applying standard linear algebra operations.

After hiding target traits in half the individuals and gene expression data in the other half, we conducted reference trait analysis and quantified performance as the fraction of the time true trait-gene expression correlations were detected using a 10% FDR threshold. *P*-values for trait-gene expression correlations were calculated using a two-sided *T* statistic and correlations deemed significant at a 10% FDR were identified using *q*-values (Storey and Tibshirani 2003).

### Imputing gene expression using TWAS

We divided the anxiety dataset in half and considered open-field measurements as target traits, hiding gene expression measurements for the animals where we did not hide open-field traits. For the TWAS strategy, our training cohort consisted of animals with genotypes and gene expression data, and our testing cohort consisted of animals with genotypes and open-field traits (i.e. training/testing labels are reversed from reference trait analysis, see Supplementary Figure 1). Diversity Outbred mice are an outbred population with genomic ancestry derived from eight inbred founder strains. We used methods implemented in R/qtl2 software (http://kbroman.org/qtl2/) to impute single nucleotide polymorphism (SNP) variation in each mouse from array-based genotypes obtained at coarser resolution (see above) using known SNP genotypes present in founder haplotypes. This resulted in genotypes for ~30 million SNPs. Given the limited number of generations of outbreeding, haplotype blocks in Diversity Outbred mice typically stretch for megabases (Svenson *et al*. 2012), leading to strong local linkage disequilibrium (LD). As such, we used PLINK version 1.9 (Purcell *et al*. 2007) to prune variants in very strong LD in the eight founder strains, using the parameters --indep-pairwise 200kb 40kb 0.95. This procedure reduced the number of SNPs to 235,335 with minimal loss of information.

To impute gene expression, we used the PredictDB module of PrediXcan (Gamazon *et al*. 2015) to build predictive models of gene expression from local genotypes within 10Mb of each gene, with sex included as a covariate. We conducted 1,000 permutations with random 50:50 divisions of the anxiety dataset to account for stochastic sampling effects. For each replicate we obtained predictive models of gene expression by running PredictDB on the training cohort and applied them to the testing cohort in order to impute gene expression. Following Gamazon et al. (2015; https://github.com/hakyimlab/PrediXcan), we considered only genes with models that were significantly predictive of gene expression (FDR ≤ 5%). Finally, we calculated correlations between imputed gene expression and a summary measure of the target traits (first canonical variable) in the testing cohort. Results were nearly identical whether we correlated to the first canonical variable or first principal component of the target traits (median transitive reliability 45% vs. 44%), but correlations to first canonical variable allow for direct comparison with results from reference trait analysis.

### Scoring gene sets to assess retrieval of known anxiety-related genes

To score gene sets for relevance to anxiety, a rater with expertise in behavioral neuroscience who was blind to the analysis methods scored a combined list of all gene sets obtained herein. We assigned a score of zero to irrelevant data sets, a score of two to gene sets with general brain or behavior relevance, and a score of four to anxiety relevant data sets in which either the gene set was generated in an anxiety relevant experiment, the gene set consisted of genes interacting with a compound known to be anxiolytic or anxiogenic, or the gene set was a Gene Ontology annotation set with direct biological relevance to anxiety. For compounds, a single MEDLINE query of the compound name and ‘anxiety’ was performed and the results of the query were examined for overall conceptual relevance.

### Data availability

Raw RNA-Seq gene expression data from the hippocampus of 258 Diversity Outbred mice are available from ArrayExpress (accession number XXX). A processed and normalized gene expression matrix is available as Supplementary Dataset 1. Phenotype data acquired via the open-field and light-dark box paradigms are available as Supplementary Datasets 2 and 3.

## Supporting information

## Acknowledgements

We thank Jackson Laboratory Genome Technologies for assistance with library preparation and sequencing of RNA-Seq samples. We thank Daniel M. Gatti for assistance with processing mouse genotypes and obtaining genotype probabilities for mapping, Juliet Ndukum for implementing an early prototype of the reference trait analysis methodology, and Timothy Reynolds for assistance with the GeneWeaver platform. This study was supported by National Institutes of Health grants R01 DA037927 (EJC), R01 AA018776 (EJC), and P50 GM076468 (EJC) and by program funds to EJC from The Jackson Laboratory.

## Author contributions

DAS – Conceptualization, Data Curation, Formal Analysis, Methodology, Software, Visualization, Writing

NR – Data Curation, Formal Analysis, Software

RFR – Investigation, Methodology

JHG – Formal Analysis

EJC – Conceptualization, Funding Acquisition, Methodology, Project Administration, Supervision, Writing

---

**Supplementary Table 1:**
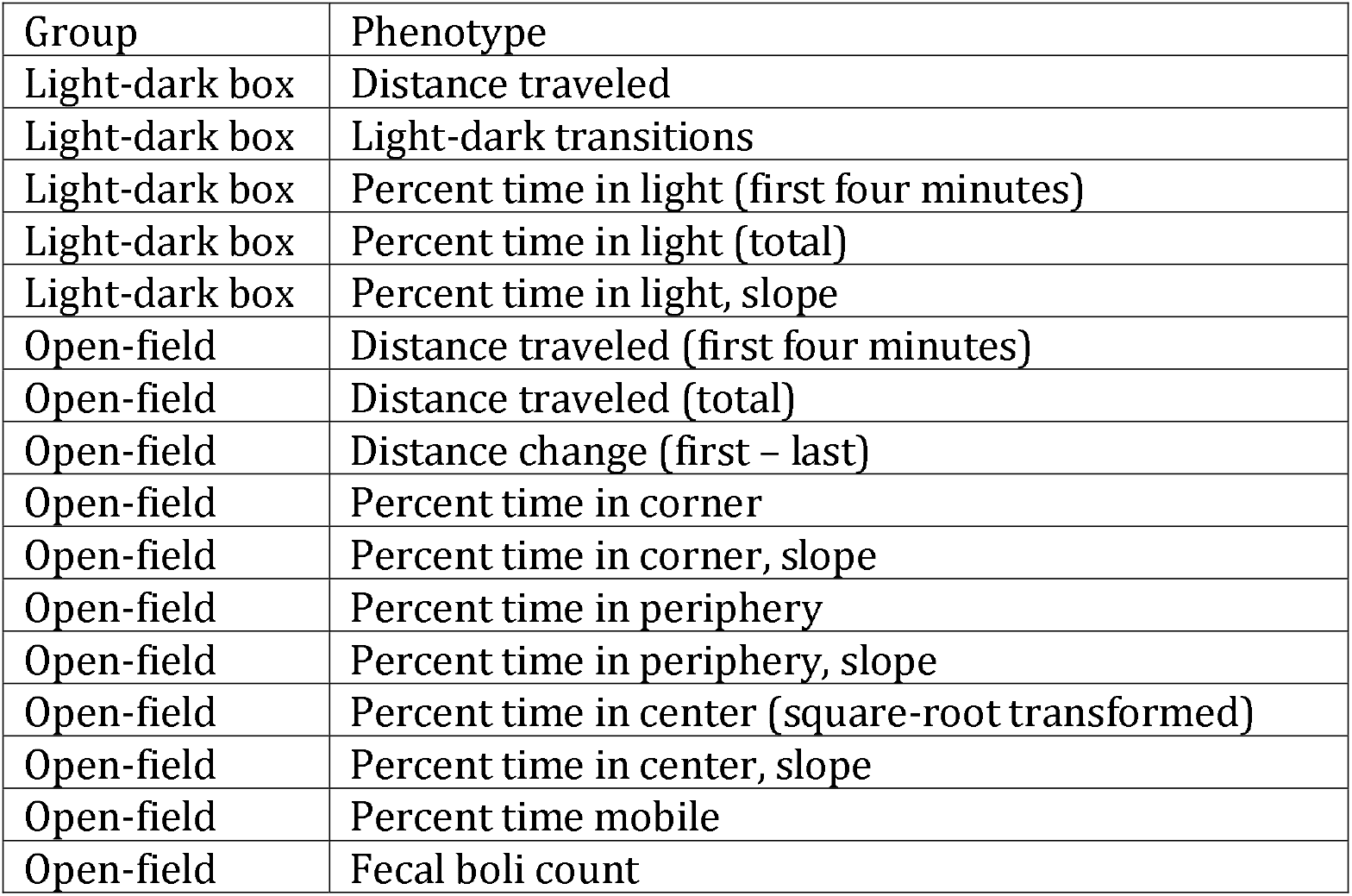
Anxiety-related traits measured on 258 Diversity Outbred mice used in case study of reference trait analysis.

| Group | Phenotype |
| --- | --- |
| Light-dark box | Distance traveled |
| Light-dark box | Light-dark transitions |
| Light-dark box | Percent time in light (first four minutes) |
| Light-dark box | Percent time in light (total) |
| Light-dark box | Percent time in light, slope |
| Open-field | Distance traveled (first four minutes) |
| Open-field | Distance traveled (total) |
| Open-field | Distance change (first – last) |
| Open-field | Percent time in corner |
| Open-field | Percent time in corner, slope |
| Open-field | Percent time in periphery |
| Open-field | Percent time in periphery, slope |
| Open-field | Percent time in center (square-root transformed) |
| Open-field | Percent time in center, slope |
| Open-field | Percent time mobile |
| Open-field | Fecal boli count |

**Supplementary Figure 1:**
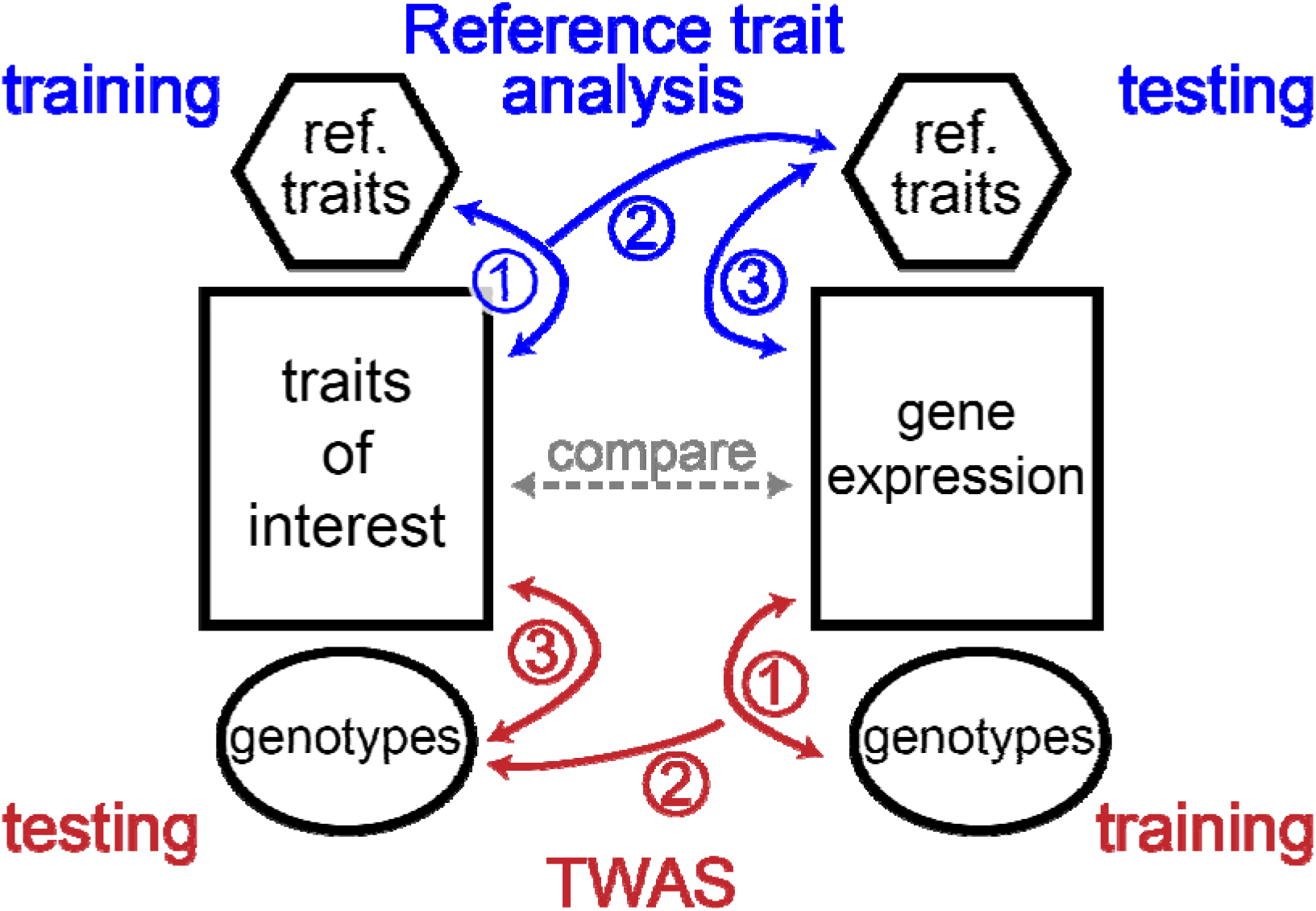
Schematic comparing overall strategies of reference trait analysis and TWAS. For reference trait analysis, canonical correlation analysis is used to relate traits of interest to reference traits (blue, 1) and coefficients derived from this model are applied to reference traits in the cohort without measurements of traits of interest (blue, 2). Finally, these projected reference traits are compared to gene expression to identify trait-gene expression correlations (blue, 3). In the TWAS approach, genotypes are used to build models that predict gene expression through eQTL (red, 1). These models are applied to genotypes in the cohort without gene expression measurements (red, 2) and imputed gene expression is compared with traits of interest to identify trait-gene expression correlations (red, 3). Note that training and testing cohort labels are switched for the two methods but that the end result of each is to compare traits of interest with gene expression (grey dashed line, middle).

**Supplementary Figure 2:**
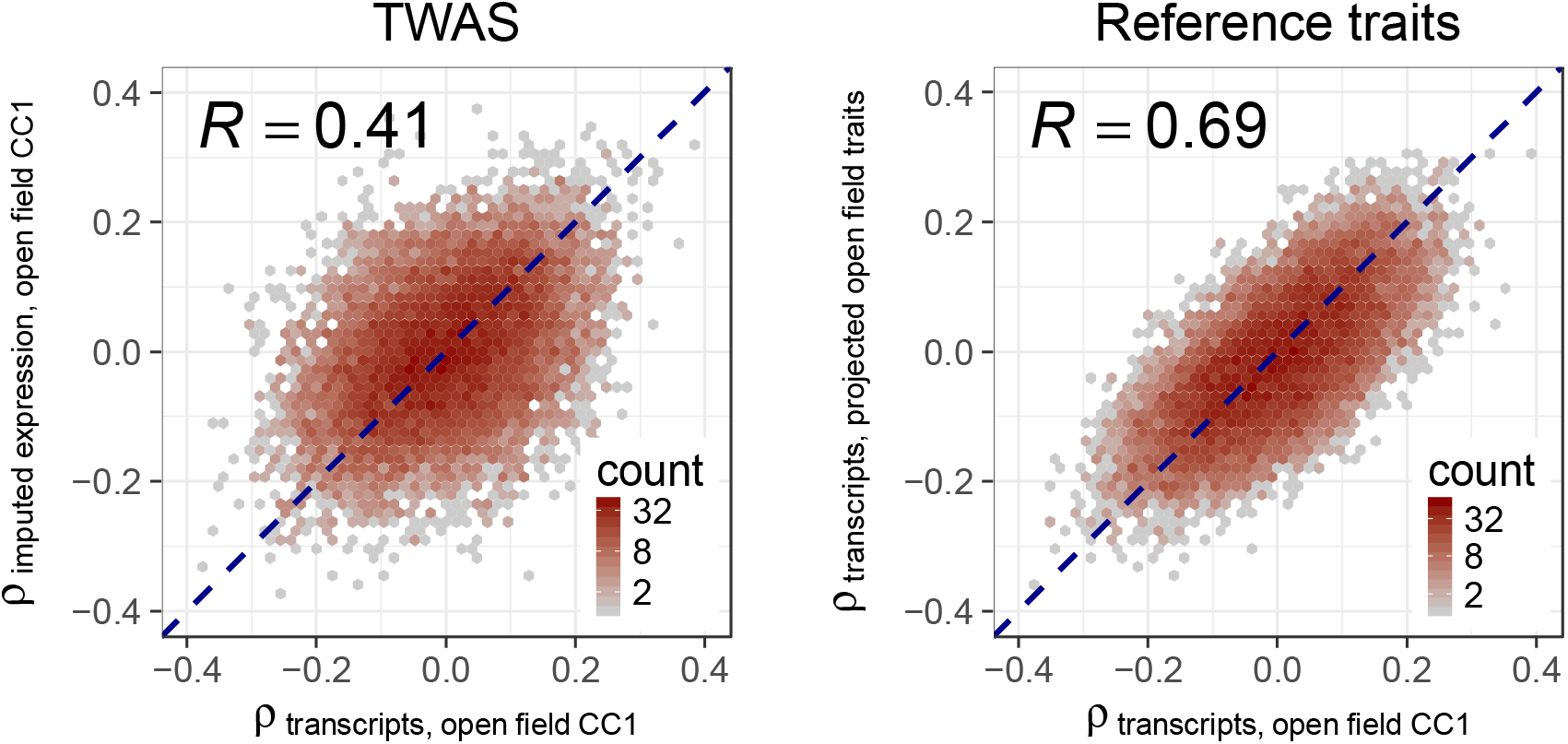
Comparison of TWAS and reference trait analysis using a single random division of the mouse anxiety dataset. For both panels we take the true trait of interest to be the first canonical covariate of open-field traits (open-field CC1). For TWAS we used genotypes to impute gene expression. Left panel shows correlation of individual transcripts to the trait of interest, where the *x*-axis plots correlations based on true transcript abundance and the *y*-axis plots correlations based on imputed transcript abundance. Right panel shows the analogous result but using reference trait analysis, where gene expression is fixed and predictors of open-field behavior are represented by projected traits.

**Supplementary Datasets**

**Supplementary Dataset 1:** Normalized hippocampal gene expression matrix. RNA-Seq data were processed as described (Methods). To obtain normalized gene expression matrix, raw counts in each sample were normalized to the upper quartile value and transformed to normal scores.

**Supplementary Dataset 2:** Traits derived from open-field arena exploration assay and used in case study of reference trait analysis. Supplementary Table 1 provides basic information on phenotypes, while complete details of animal rearing, husbandry and phenotyping are presented in Logan et al. (2013).

**Supplementary Dataset 3:** Traits derived from light-dark box behavior assay and used in case study of reference trait analysis. Supplementary Table 1 provides basic information on phenotypes, while complete details of animal rearing, husbandry and phenotyping are presented in Logan et al. (2013).

